# It’s just a phase: moonlight and rainfall influence the activity of a threatened cave-roosting bat in the Pilbara

**DOI:** 10.64898/2026.08.20.746117

**Authors:** E. L. Westerhuis, M. Kaestli, C. Grabham, H. North, K. Armstrong, G. Madani, J. O’Brien

## Abstract

Wildlife monitoring in arid environments is complicated by substantial temporal and spatial variation in animal activity driven by unpredictable environmental conditions. For highly mobile species, movements across the landscape may further influence activity recorded at fixed monitoring locations, raising questions about whether short-term surveys adequately represent patterns of site use. We examined temporal variation in acoustic activity of the threatened Pilbara Leaf-nosed Bat (*Rhinonicteris aurantia* Pilbara form), a highly mobile, cave-roosting insectivorous bat inhabiting the arid Pilbara region of north-western Australia. Acoustic activity was monitored continuously at two permanent diurnal roosts for up to two years, comprising 1,601 detector-nights across three detector locations. We tested relationships between nightly activity and environmental conditions, detector location and anthropogenic disturbance. Activity varied substantially through time and among detector locations. Moon illumination and cumulative rainfall were strongly associated with activity, but their interaction differed between roosts: activity was highest under low moon illumination and low cumulative rainfall at Chateau Cave, whereas activity at Daltons was highest under low moon illumination and higher cumulative rainfall. Activity also differed markedly between detector positions within Chateau Cave, and humidity, temperature and artificial light at night had additional location-specific effects. Other measures of anthropogenic disturbance, including cave entry, blasting and mining activity, had comparatively weak support. The contrasting environmental relationships among locations demonstrate that acoustic activity at permanent roosts is strongly context dependent. For a species capable of extensive movements among roosts, temporal variation in activity may reflect behavioural responses and redistribution of individuals across the landscape rather than rapid demographic change. We suggest that facultative nomadism may provide a useful hypothesis for understanding temporal variation in roost use by Pilbara Leaf-nosed Bat. More broadly, our results demonstrate that short-duration acoustic surveys of highly mobile species in arid environments may provide an incomplete representation of longer-term site use and should incorporate temporal replication and environmental context when used for conservation and impact assessment.

## Introduction

Arid ecosystems are driven by highly variable and unpredictable rainfall with high evapotranspiration, resulting in pronounced temporal and spatial variation in resource availability (Noy-Meir 1973; Morton et al. 2011). Arid-adapted plants and animals exhibit a wide range of physiological, behavioural and life-history adaptations to these conditions, including physiological tolerance, dormancy, rapid reproduction following resource pulses, selection of favourable microhabitats and movement to track spatially variable resources (Ward 2016).

Mammals exhibit a diverse suite of physiological, behavioural and life-history adaptations to arid environments and encompass a wide range of body sizes, life histories and reproductive strategies, from small rodents, bats and marsupials to large herbivores and carnivores (Lewin et al. 2024). These differences influence how species respond to spatial and temporal variation in resources. Populations of small-bodied, short-lived terrestrial mammals, particularly rodents, can respond to rainfall-driven increases in resource availability through rapid reproduction and population growth, followed by declines as conditions deteriorate (Ostfeld and Keesing 2000; Dickman et al. 1999; Dickman et al. 2011). Large-bodied mammals with slower reproductive rates may instead respond to resource variability through extensive movements that track spatially dispersed resources (Nandintsetseg et al. 2019). Where resources are particularly unpredictable in space and time, such movements may take the form of nomadism, a strategy widely associated with animals inhabiting variable and unpredictable environments (Teitelbaum and Mueller 2019).

Bats occupy an unusual position within this spectrum, combining small body size with comparatively long lifespans and low reproductive rates, while powered flight provides exceptional mobility. Despite the energetic and water-balance challenges associated with flight, bats are a diverse and successful component of arid mammal communities (Conenna et al. 2025), second only to rodents, with at least 248 species represented among predominantly dryland mammals (Lewin et al. 2024). Movement is an important behavioural response of bats to resource limitation in arid environments, occurring over timescales ranging from individual nights to entire seasons (Conenna et al. 2025). Insectivorous bats may commute long distances to access water and foraging resources, expand their foraging ranges during dry periods, undertake seasonal movements, or move to alternative roosts as resources vary spatially and temporally (Conenna et al. 2025). Consequently, the presence or activity of bats at a particular site may be influenced not only by conditions at that site, but also by the distribution and availability of resources across the surrounding landscape. Temporal changes in local activity may therefore reflect redistribution of individuals across the landscape rather than equivalent changes in population abundance.

Cave-roosting insectivorous bats in arid environments provide an ideal system for examining these dynamics because they depend on spatially fixed and temporally persistent roost refugia while exploiting food resources that vary substantially across the surrounding landscape (Furey and Racey 2016). Invertebrate abundance in arid ecosystems is influenced by both vegetation structure and rainfall, but taxa differ in the magnitude and timing of their responses to rainfall, resulting in highly variable and often unpredictable patterns of prey availability (Palmer 2010; Kwok et al. 2016; Maute et al. 2019). Consequently, conditions at a roost may remain relatively stable while the resources available to bats using that roost change substantially through space and time.

Among Australia’s relatively small number of arid-zone cave-roosting bats, the Pilbara Leaf-nosed Bat (*Rhinonicteris aurantia* Pilbara form; syn. Pilbara diamond-faced bat, Armstrong et al. 2016) provides an especially informative example because its distribution is restricted to one of the continent’s most climatically variable arid regions and it has highly specialised roost requirements. The Pilbara Leaf-nosed Bat is recognised as a separate form and isolated population of the more widespread Orange Leaf-nosed Bat (*Rhinonicteris aurantia*, Grey 1845). *Rhinonicteris aurantia* is the only surviving Australian species of the Trident Bat Family (*Rhinonycteridae*), a small family of otherwise paleotropical species (Armstrong 2023). The species has one of the highest rates of pulmocutaneous water loss for any mammal (Baudinette et al. 2020). Because individuals are unable to passively maintain body temperature at ambient temperatures below 30 °C (Baudinette et al. 2020), they have obligate requirements for warm, humid roost microclimates (Armstrong 2000; Armstrong 2001).

The persistence of a physiologically delicate cave-roosting bat in one of Australia’s hottest and most climatically variable regions presents an intriguing ecological paradox. In the Pilbara, daily maximum temperatures greater than 45 °C are not uncommon and several instances of temperatures exceeding 50 °C have been recorded. In addition to high temperatures, the Pilbara, like much of arid Australia, is characterised by high spatial and temporal rainfall variability (Morton et al. 2011). Average rainfall varies from 250 to 350 mm per year, but interannual rainfall variability is high, second only to central Australia (Van Etten et al. 2009). Compounding these climatic challenges, the Pilbara is one of Australia’s most intensively mined regions, and high-grade iron ore deposits frequently occur within the same banded ironstone formations that support critical roost habitat for the Pilbara Leaf-nosed Bat (Bat Call WA 2021; Cramer et al. 2016).

Records of Pilbara Leaf-nosed Bat are distributed throughout the Pilbara region but are concentrated in rocky areas that provide roosting opportunity (Armstrong 2001, 2003). They are easily identified from acoustic recordings, and survey effort has to date been most extensive in areas where mining occurs (Bradley et al. 2024). There are approximately 60 roosts known or suspected (Shaw et al. 2026), supporting an estimated population of 10,000–22,000 individuals (Bat Call WA 2021). The loss of diurnal roosts is a key threatening process (Threatened Species Scientific Committee 2016), while predation from both native and non-native predators (Moyses et al. 2024) and land-use change are also recognised threats (Cramer et al. 2016; Bradley et al. 2024).

Despite being physiologically sensitive and reliant on stable diurnal roosts, Pilbara Leaf-nosed Bat are capable of extensive movements. One male was recorded travelling 170 km between diurnal roosts over several months (Bullen and Reiffer 2019), and individuals have been documented undertaking nightly foraging flights of up to 40 km (Knuckey et al. 2024). More recently, females have been recorded moving between roosts up to 60 km apart within a few hours and exhibiting frequent roost-switching behaviour over consecutive nights (O’Brien and Westerhuis 2025). These extensive and apparently irregular movements raise the possibility that Pilbara Leaf-nosed Bat may exhibit facultative nomadism, whereby individuals shift their spatial distribution in response to resources or environmental conditions that vary unpredictably across the landscape (Teitelbaum and Mueller 2019). This interpretation is further supported by high rates of dispersal and limited genetic population structure (Umbrello et al. 2022), consistent with landscape-level fission-fusion social dynamics (Kerth and König 1999). Consequently, the number of bats using an individual diurnal roost may vary through time as individuals move among roosts, meaning observations made during a short survey period may not necessarily reflect the longer-term importance of that roost.

Understanding the environmental drivers of temporal variation in activity is therefore important for interpreting patterns of roost use. Pilbara Leaf-nosed Bat activity has been hypothesised to be driven primarily by annual rainfall or seasonal transitions between the wet and dry seasons (Bat Call WA 2021), although this has not been formally tested. More broadly, bat activity is known to vary in response to temperature (Koch et al. 2023; Turbill 2008; Meyer et al. 2004), rainfall (Milne et al. 2005), humidity (Lacki 1984; Rodríguez-San Pedro et al. 2024), wind speed (Arnett and Baerwald 2013), barometric pressure (Bender and Hartman 2015; Paige 1995), lunar phase (Börk 2006; Zeppelini et al. 2019; Vasquez et al. 2020), invertebrate abundance (Bhalla et al. 2023) and fire (Doty et al. 2016; Law et al. 2018).

Acoustic monitoring is widely used to survey Pilbara Leaf-nosed Bat because it provides a relatively inexpensive and effective means of collecting data. However, the number of acoustic detections recorded at a site represents bat activity rather than the number of individuals present. A single individual may generate multiple detections, while variation in flight behaviour, foraging activity and movement past a detector can alter the number of calls recorded independently of abundance (Gorresen et al. 2008; Udell et al. 2024). The period over which acoustic monitoring is conducted may therefore strongly influence conclusions about the relative importance of a site.

Together, high mobility, environmental variability and the indirect relationship between acoustic activity and abundance create substantial uncertainty when interpreting short-term monitoring at diurnal roosts. Quantifying the magnitude and drivers of temporal variation in activity is therefore necessary to determine whether short-term acoustic surveys provide representative measures of roost use and to distinguish natural variation from responses to anthropogenic disturbance.

We used 24 months of continuous acoustic monitoring at two diurnal roosts of the Pilbara Leaf-nosed Bat to investigate how natural environmental variability and human disturbance shape activity patterns in one of Australia’s most climatically unpredictable arid regions. Specifically, we asked:

1. How much does Pilbara Leaf-nosed Bat activity vary among roosts, detector positions and through time?
2. Which environmental variables are associated with temporal variation in acoustic activity?
3. Is acoustic activity associated with anthropogenic disturbance, including artificial light at night, cave entry, blasting, mining activity and fire?

We then consider the implications of temporal variation for interpreting acoustic surveys and designing monitoring programs for threatened cave-roosting bats in climatically variable landscapes.

## Methods

### Study area and data collection

Two diurnal roosts in natural caves were chosen for this study. Chateau Cave is a relatively deep (∼20 m) cave within banded ironstone. The cave is located within the North Star Magnetite Project development envelope, 115 km southeast of Port Hedland in the Chichester region of the Pilbara, and operated by FMG Iron Bridge Pty. Ltd. A 100 m mining exclusion zone was established around the roost as a condition of the Project’s environmental approvals. The cave entrance is near the top of an escarpment, with a narrow passageway leading to a chamber where Pilbara Leaf-nosed Bat roost. The roost area is completely dark. Two full-spectrum ultrasonic detectors (Song Meter SM4BAT-FS, Wildlife Acoustics) were placed at Chateau Cave in early April 2021. One was placed inside the internal roosting chamber of the cave (hereafter ‘Chateau Cave in’) and paired with a second detector positioned outside the cave, 5 m from the entrance (Chateau Cave out’).

A concurrent genetic mark-recapture study and hormone analysis was conducted using Pilbara Leaf-nosed Bat scat collected from beneath the main roosting area at Chateau Cave. This involved cave entry once per month for two consecutive nights (Armstrong et al. in prep, Westerhuis et al. in prep). Analysis of scats and thermal camera counts indicates that the Chateau colony consists of approximately 1500 individuals, with a high degree of individual turnover (Armstrong et al. in prep). Sex-linked markers have also shown that the colony has an overall male bias, although this varies month to month. The role of the cave in the breeding cycle remains unclear. Hormone analysis of scats has shown that pregnant females regularly roost at this location during later stages of the presumed pregnancy period (Westerhuis et al. in prep), but observations of pups in the cave have been limited to two pups on one night in February 2022, despite regular inspections.

A second diurnal roost, Daltons Creek Roost (hereafter ‘Daltons’), was added to the study in April 2022, with data from June 2021 to December 2021 provided by Biologic Environmental Consultancy and Atlas Iron Pty Ltd. Daltons is a cave in conglomerate rock 29 km southeast of Chateau Cave (Figure 1). All Daltons monitoring used SM4BAT-FS detectors, but different units were used by Biologic and the GHD team. All units were scheduled to record from 30 minutes before sunset through to 30 minutes after sunrise each night. The nearest active mining operation to this roost is Mt Webber, operated by Atlas Iron, located more than 10 km southeast of the cave. Scat collection began at this roost in May 2022 but did not occur every month and, due to physical access limitations, was only undertaken outside the cave entrance. In contrast to Chateau Cave, the sex bias at Daltons is female-skewed (Armstrong et al. in prep). Hormone analysis of scats also shows pregnant females are present at this location in later stages of the pregnancy period (Westerhuis et al. in prep), and females trapped at the site in December 2022 were all heavily pregnant. Cat predation has been noted regularly at Daltons (Moyses et al. 2024), although monitoring of cat predation has not been continuous. Before GHD ecologists commenced monitoring, a wildfire burnt the area surrounding Daltons in November 2021. This fire damaged the Biologic bat detector and consequently no data were recorded for January–March 2022 until the GHD detector was installed. Acoustic detectors were programmed to record nightly throughout the study period, providing near-continuous monitoring except during periods of equipment failure or other technical difficulties.

**Figure 1.**
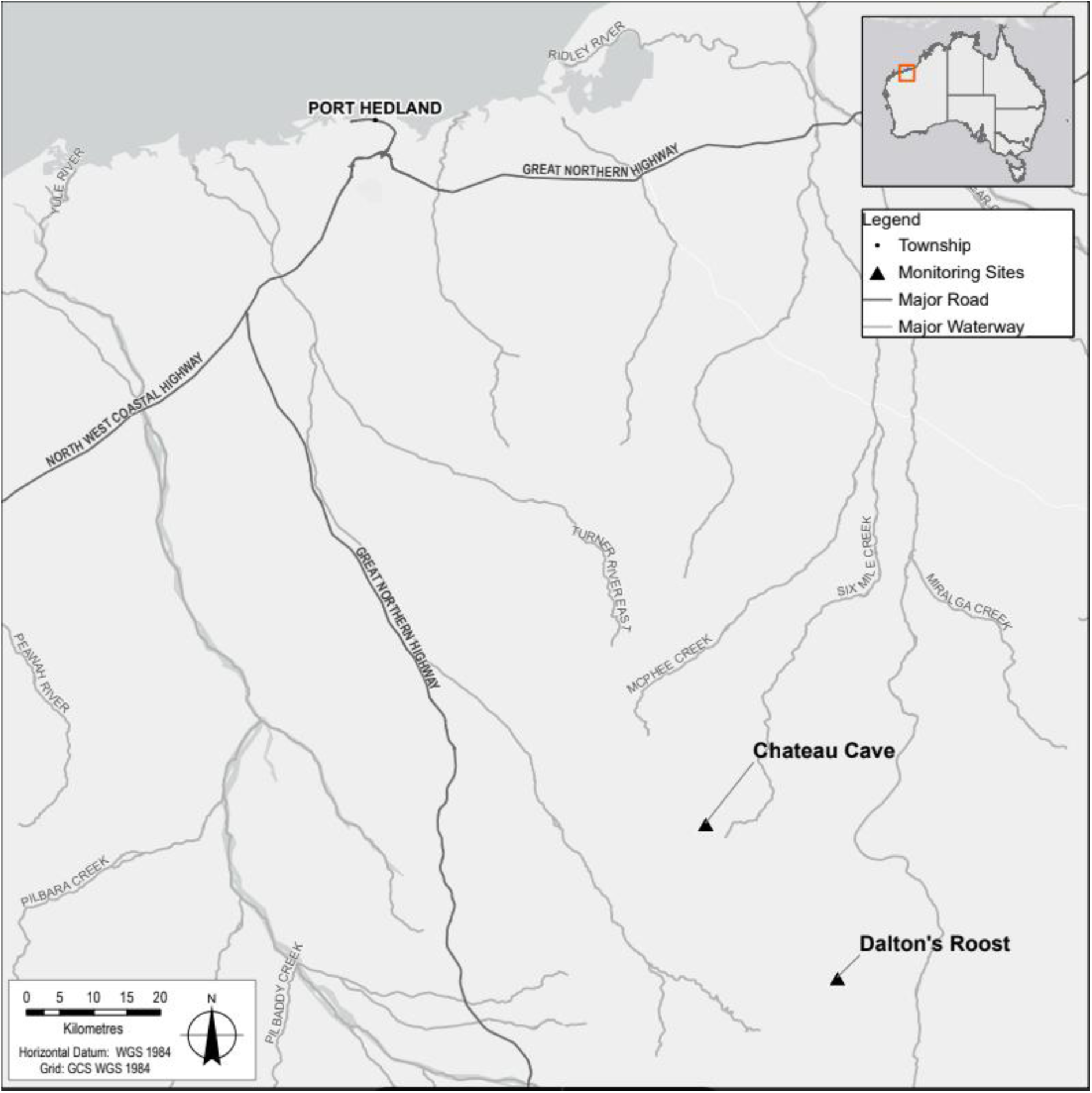
Location of the two roosts used in the study.

All GHD monitoring was conducted under “Licence to Take Fauna for Scientific Purposes” (Regulation 17) Permit No. 08-000751-1 and Animal Research Authority (Project number GHD: 613519501 and 12548779).

### Call identification and data processing

Data were processed and analysed using a combination of manual review and automated processes in Kaleidoscope Pro (Wildlife Acoustics, version 5.1.9) and Anabat Insight (Titley Scientific, version 1.9.9-7). Files were split into 1 s lengths as the basic unit for summing bat call activity. Data for each night and site were processed using the Kaleidoscope Pro cluster analysis function and a species-specific classifier for Pilbara Leaf-nosed Bat developed from a manually vetted training dataset of Pilbara Leaf-nosed Bat files from the study area. The training dataset consisted of 5500 files, of which 5000 contained Pilbara Leaf-nosed Bat calls and 500 contained calls of other species. Following refinement, the classifier returned a 98% true positive rate, with the remaining 2% of files comprising false positives (1.2% of calls incorrectly identified as other species) and false negatives (0.8% missed Pilbara Leaf-nosed Bat calls). False positive files were removed before data analysis. At least one clear pulse was accepted because Pilbara Leaf-nosed Bat echolocation calls are distinctive, with constant frequency structure and a characteristic frequency of 116–126 kHz (Armstrong and Coles 2007). Acoustic activity was interpreted as a relative index of recorded activity rather than a direct count of bats.

### Environmental variables

Concurrent with the acoustic monitoring period, weather data comprising temperature, humidity, barometric pressure, rainfall and wind speed were downloaded for each 15-minute interval from the Iron Bridge weather station. Minimum, maximum and mean values were calculated for each day. Moon illumination for each night was calculated using the Suncalc package (Thieurmel & Elmarhraoui 2022) and the GPS coordinates for Chateau Cave.

FMG provided data on blasting and mining activity, including distance from Chateau Cave to the blast site, blast depth and vibration measured at the cave. Distance to blast was used to create a binary factor of before and after mining based on increased mining activity after 17 November 2021, when mining pits near the cave became operational. The artificial light factor was created based on whether a light tower illuminating mining infrastructure was operational. The light operated intermittently in late 2022 and continuously from early 2023. A lux meter, along with noise and vibration loggers, was deployed; however, technical issues resulted in substantial missing data and these data were not used further. The binary fire factor was created based on dates before and after 22 November 2021. The binary cave-entry factor was coded as yes for each night cave entry occurred and for three days after entry. Although cave entry was always conducted after civil twilight, when Pilbara Leaf-nosed Bat were anticipated to have emerged from the cave, a lag effect was considered plausible given the known sensitivity of cave-roosting bats to roost disturbance.

We checked for collinearity among continuous predictor variables using pairwise Spearman’s rank correlation coefficients. Paired predictors with correlation coefficients exceeding 0.7 were considered highly correlated, and only the predictor with the strongest association with Pilbara Leaf-nosed Bat activity was retained for further analysis. This approach was used to reduce redundancy among predictors and improve interpretability of model outputs.

### Statistical analysis

#### 1. Temporal and spatial variation in Pilbara Leaf-nosed Bat activity

To quantify temporal and spatial variation in Pilbara Leaf-nosed Bat activity, we summarised nightly activity across the three detector locations (Chateau Cave in, Chateau Cave out and Daltons) and months from April 2021 to March 2023. We calculated descriptive measures of the magnitude and distribution of nightly activity within each detector location and survey period and used a univariate permutational analysis of variance (PERMANOVA) in PRIMER-e (Quest Research, Auckland, New Zealand) with the PERMANOVA+ add-on package to test for broad differences in activity among detector locations and survey periods. PERMANOVA was applied to log-transformed activity because it can accommodate complex unbalanced designs and non-normal data (Anderson et al. 2008). Because nightly observations were temporally structured, these results were interpreted as evidence of broad spatial and temporal differences in activity rather than as a mechanistic analysis of temporal drivers. PERMDISP was also used to assess differences in dispersion among detector locations and survey periods.

Due to high variance among sites and strong right skew in the bat activity data, activity was log-transformed before analysis. A Euclidean distance similarity matrix was then calculated based on the transformed data. Euclidean distance was chosen because it is appropriate for univariate analysis and, when used in PERMDISP, is equivalent to a Levene’s test of dispersion (Anderson et al. 2008).

Both monthly survey (n=25, April 2021–April 2023) and detector location (n=3, Chateau Cave in, Chateau Cave out and Daltons) were treated as fixed factors. Site was treated as fixed because the two diurnal roosts in this study represent a defined subset of known diurnal roosts and the objective was to compare those specific roosts and detector positions rather than extrapolate to all Pilbara Leaf-nosed Bat roosts. Survey period was also treated as fixed because the analysis aimed to compare activity among specific survey periods. The design was unbalanced, so 9,999 permutations with unrestricted permutations of raw data were used to estimate p-values. Type III sums of squares were used to account for the unbalanced design, and fixed effects were set to sum to zero for mixed terms.

#### 2. Models of Pilbara Leaf-nosed Bat activity against environmental variables

Nightly activity patterns at both diurnal roosts were analysed using the nightly sum of acoustic activity from 18:00 to 06:00. Rows with missing values for relevant environmental predictors, anthropogenic variables and temporal grouping variables were removed. A summary of variables is provided in Table 1.

**Table 1.**
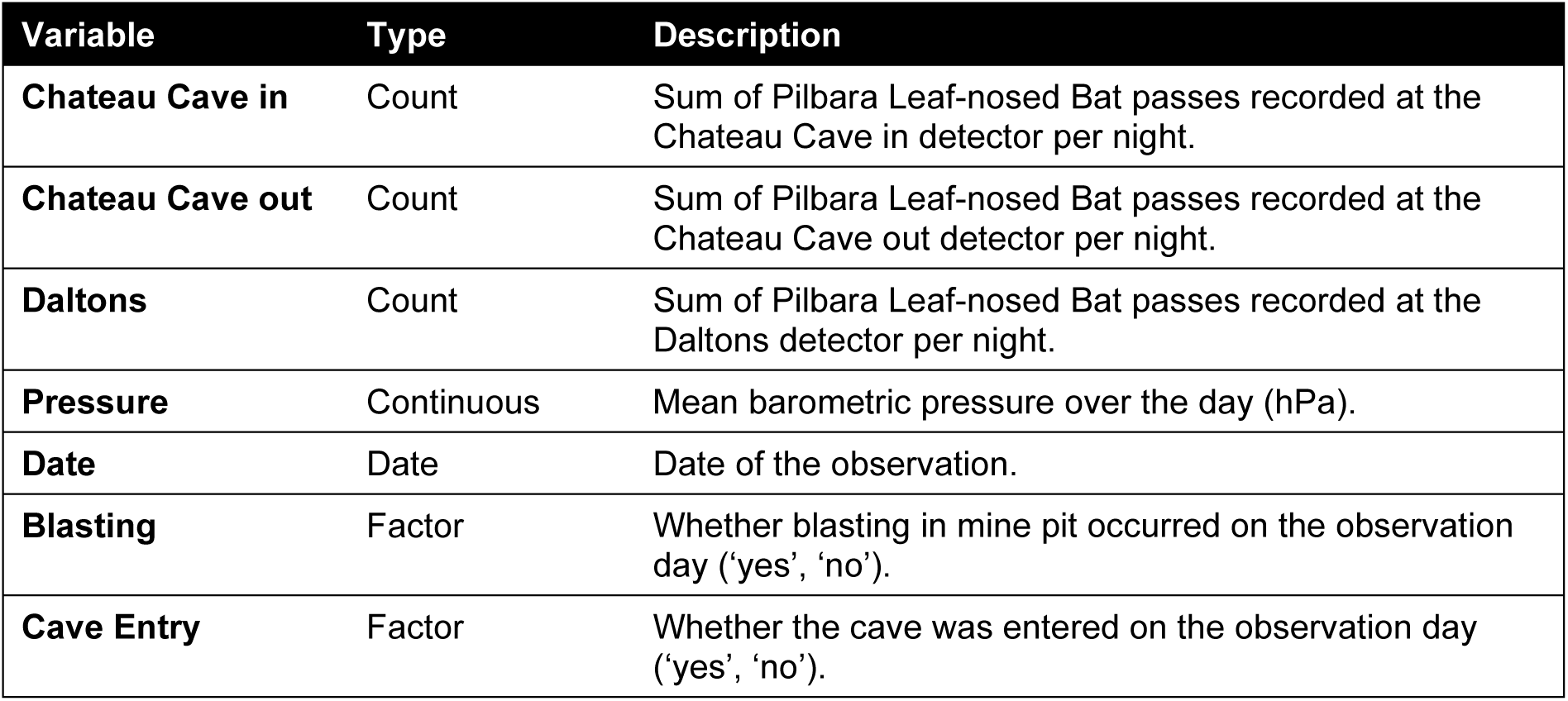
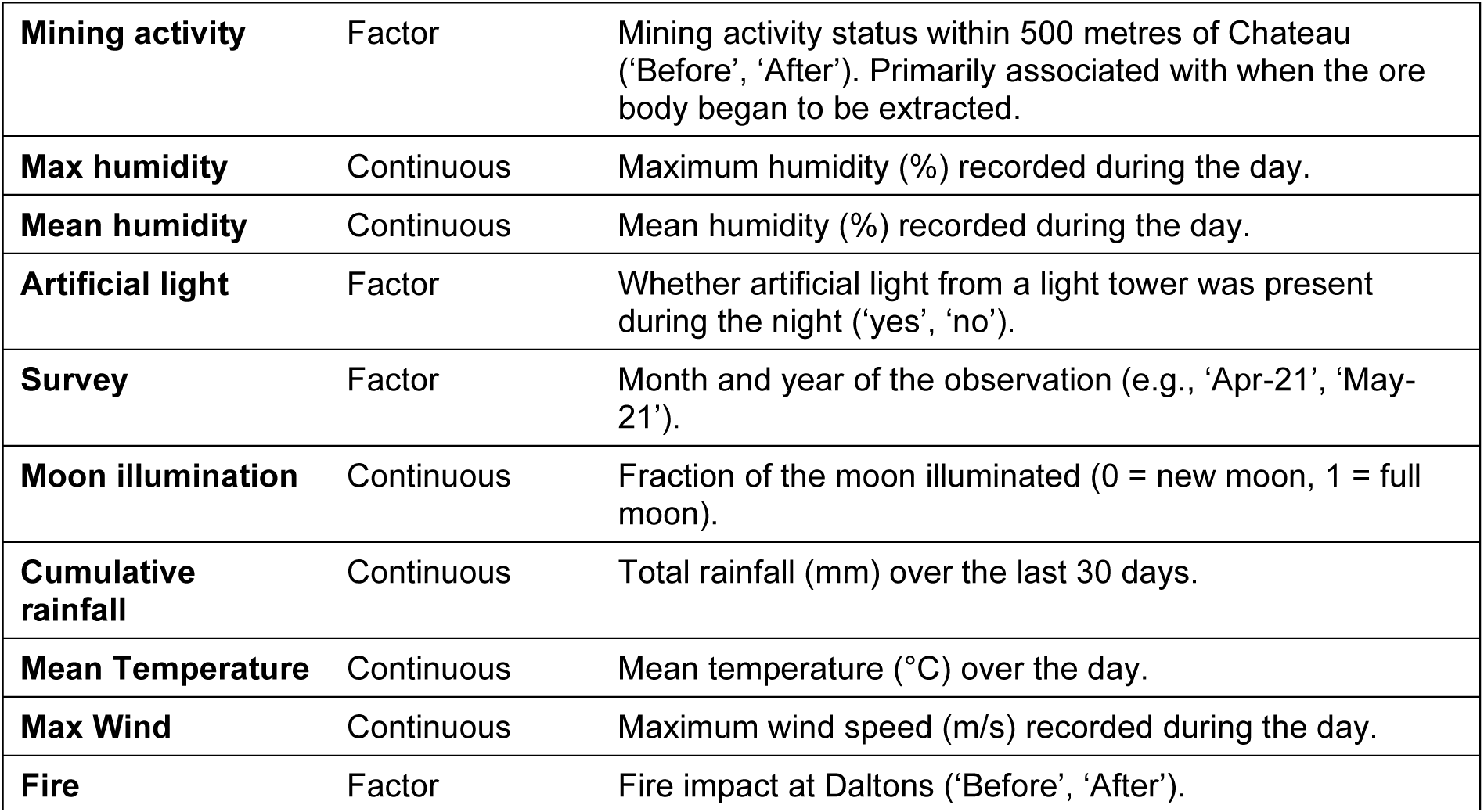
Variables used in generalised additive models.

| Variable | Type | Description |
| --- | --- | --- |
| **Chateau Cave in** | Count | Sum of Pilbara Leaf-nosed Bat passes recorded at the Chateau Cave in detector per night. |
| **Chateau Cave out** | Count | Sum of Pilbara Leaf-nosed Bat passes recorded at the Chateau Cave out detector per night. |
| **Daltons** | Count | Sum of Pilbara Leaf-nosed Bat passes recorded at the Daltons detector per night. |
| **Pressure** | Continuous | Mean barometric pressure over the day (hPa). |
| **Date** | Date | Date of the observation. |
| **Blasting** | Factor | Whether blasting in mine pit occurred on the observation day ('yes', 'no'). |
| **Cave Entry** | Factor | Whether the cave was entered on the observation day ('yes', 'no'). |
| **Mining activity** | Factor | Mining activity status within 500 metres of Chateau ('Before', 'After'). Primarily associated with when the ore body began to be extracted. |
| **Max humidity** | Continuous | Maximum humidity (%) recorded during the day. |
| **Mean humidity** | Continuous | Mean humidity (%) recorded during the day. |
| **Artificial light** | Factor | Whether artificial light from a light tower was present during the night ('yes', 'no'). |
| **Survey** | Factor | Month and year of the observation (e.g., 'Apr-21', 'May-21'). |
| **Moon illumination** | Continuous | Fraction of the moon illuminated (0 = new moon, 1 = full moon). |
| **Cumulative rainfall** | Continuous | Total rainfall (mm) over the last 30 days. |
| **Mean Temperature** | Continuous | Mean temperature (°C) over the day. |
| **Max Wind** | Continuous | Maximum wind speed (m/s) recorded during the day. |
| **Fire** | Factor | Fire impact at Daltons ('Before', 'After'). |

Generalised additive models (GAMs) were used to assess drivers of bat activity and account for non-linear relationships. Models were fitted using bam() in the mgcv package (Wood 2003, Wood 2004, Wood 2011, Wood et al. 2016, Wood 2017) in R. Although the response variable was derived from counts of one-second call files, nightly totals were large, non-zero and right-skewed. A Gamma distribution with a log link provided the best fit for Chateau Cave in and Chateau Cave out activity based on residual diagnostics using the DHARMa package (Hartig 2022). For Daltons, a Gaussian distribution with identity link provided the best fit and no negative activity was predicted. Default fast REML computation was used for model fitting, with discretisation of covariates (discrete=TRUE) for the Gamma models.

An autoregressive structure (AR1) was included to model temporal dependence in daily activity data. The AR1 correlation parameter was estimated using the start_value_rho() function in the itsadug package (van Rij et al. 2022) with the default lag-2 setting. Default thin-plate regression splines were fitted for continuous predictors and tensor product smooths were used for interactions between covariates. The base and all candidate models included a cyclic cubic spline smooth for day of year (DOY) to account for seasonal patterns not explained by environmental variables, and a categorical variable for year to capture interannual differences.

Multiple candidate models were developed to test combinations of disturbance variables, including cave entry and mining activity, and environmental predictors, including rainfall, temperature, moon illumination and wind speed. Relative model support was assessed using Akaike’s Information Criterion corrected for small sample size (AICc; Burnham and Anderson 2002) calculated within the mgcv package. Model residuals were checked for random patterns across fitted values, predictors and time using the DHARMa package, itsadug diagnostic plots and gam.check() in mgcv. Remaining temporal autocorrelation was assessed using autocorrelation function plots. Data processing was conducted using dplyr (Wickham et al. 2023), while broom (Robinson et al. 2024), ggplot2 (Wickham 2016), visreg (Breheny and Burchett 2017) and itsadug were used to summarise and visualise model outputs. All GAMs and associated data processing were conducted in R Statistical Software (version 4.4.1 “Race for Your Life”, R Core Team 2024).

## Results

### Spatial and temporal variation in nightly Pilbara Leaf-nosed Bat activity

Across the study, 1601 detector-nights were recorded across the three detector detector locations: Chateau Cave in, Chateau Cave out and Daltons. Across 609 days of recording, mean Pilbara Leaf-nosed Bat activity at the Chateau Cave out detector was 369 (±14 SE) passes per night. Activity outside Chateau Cave was generally lower than inside the roosting chamber, although 10.5% of nights recorded extremely low activity (<50 Pilbara Leaf-nosed Bat passes per night), mostly during February and March 2023. Activity inside Chateau Cave was approximately 22 times higher, with an average of 8139 (±246 SE) passes per night. Lowest activity at this detector also occurred in February 2022, February 2023 and March 2023. The average activity at Daltons was 12040 (±262 SE) passes per night across 464 detector-nights.

PERMANOVA indicated significant differences in activity among detector locations, survey periods and the detector location × survey interaction (Table 2, Figure 2). PERMDISP also indicated significant dispersion for both detector location and survey period. Pairwise PERMANOVA results showed that significant differences were more common than non-significant differences, indicating high temporal and spatial variation in activity. Differences were particularly evident between Chateau Cave in and Chateau Cave out, indicating that activity outside the cave entrance was not representative of activity inside the roosting chamber. Non-significant comparisons occurred primarily between Chateau Cave in and Daltons and in some month-specific comparisons, such as June 2021 and July 2021. Dispersion was heterogeneous for both Chateau Cave detectors, while Daltons showed greater homogeneity and more uniform activity. Overall, Pilbara Leaf-nosed Bat activity varied strongly among detector detector locations and survey periods, indicating that short-term acoustic activity may not provide a stable measure of roost use.

**Table 2.**
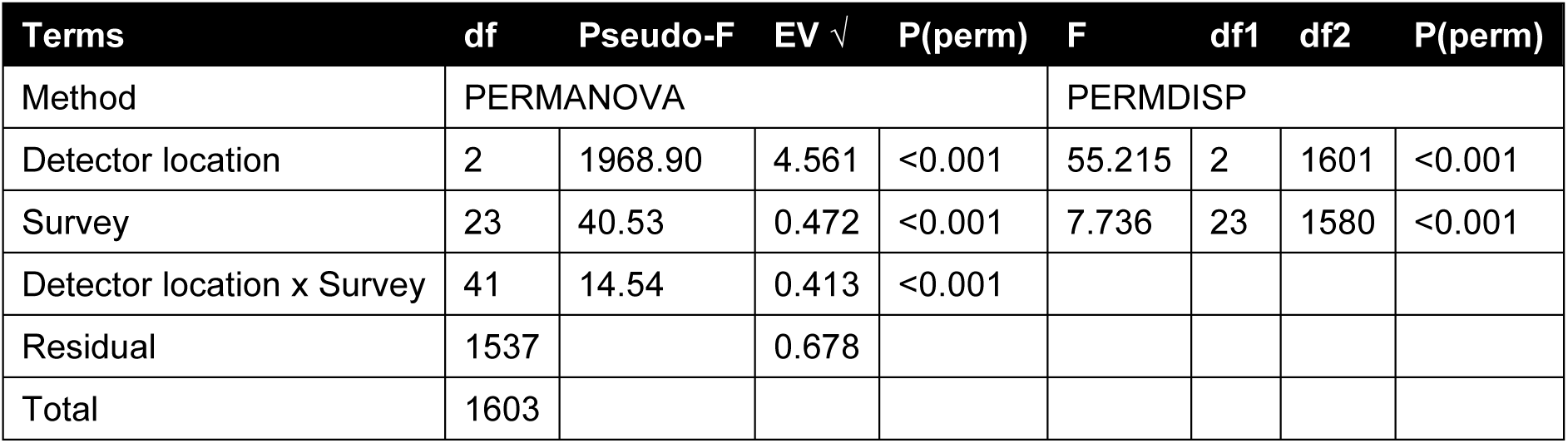
Significant univariate 2 Factor PERMANOVA on Pilbara Leaf-nosed Bat activity. Df = degrees of freedom, SS = sum of squares, EV = estimated variance, perms = permutations.

| Terms | df | Pseudo-F | EV $\sqrt{\phantom{x}}$ | P(perm) | F | df1 | df2 | P(perm) |
| --- | --- | --- | --- | --- | --- | --- | --- | --- |
| Method | PERMANOVA |  |  |  | PERMDISP |  |  |  |
| Detector location | 2 | 1968.90 | 4.561 | <0.001 | 55.215 | 2 | 1601 | <0.001 |
| Survey | 23 | 40.53 | 0.472 | <0.001 | 7.736 | 23 | 1580 | <0.001 |
| Detector location x Survey | 41 | 14.54 | 0.413 | <0.001 |  |  |  |  |
| Residual | 1537 |  | 0.678 |  |  |  |  |  |
| Total | 1603 |  |  |  |  |  |  |  |

**Figure 2.**
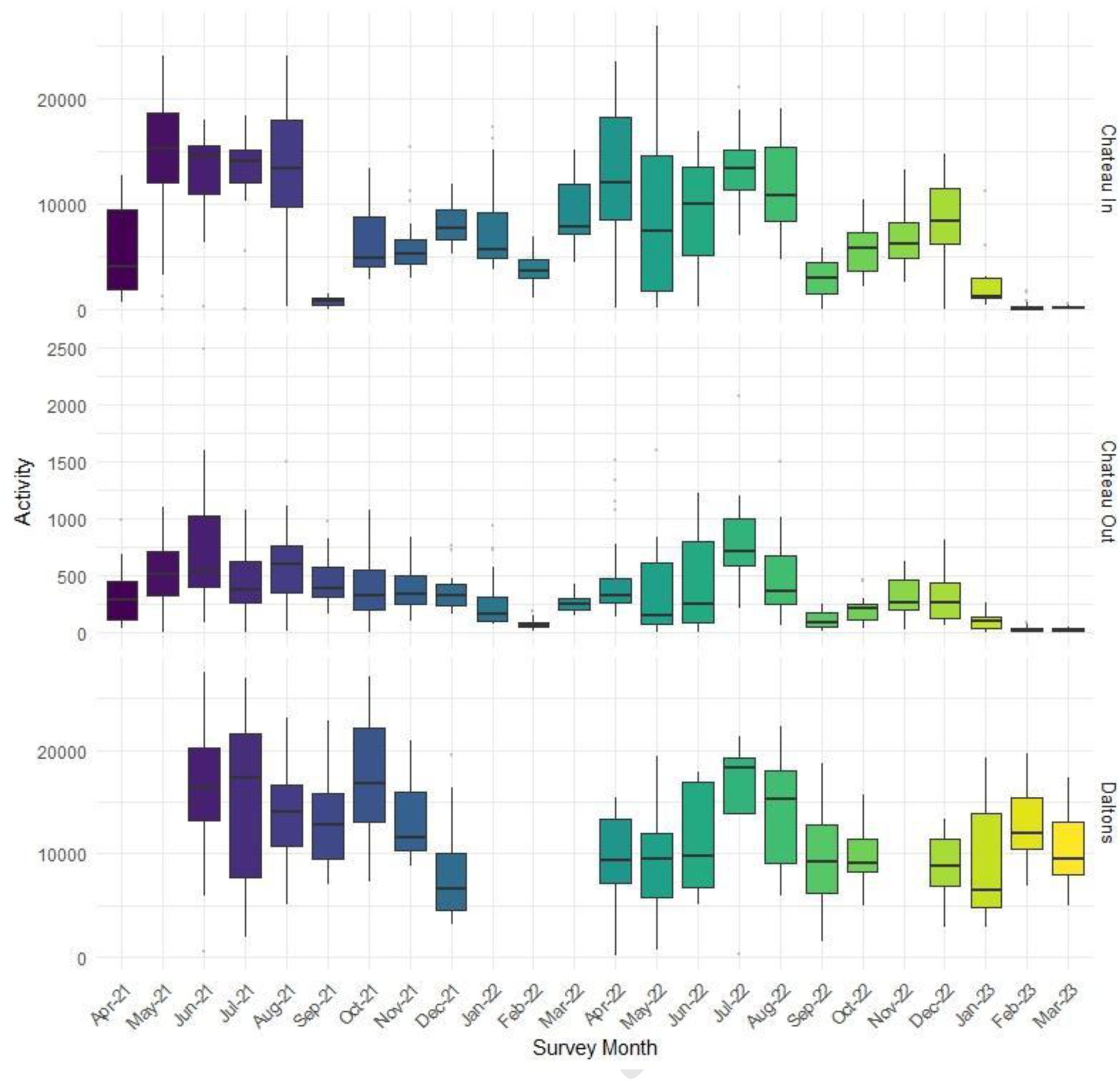
activity per night between detector locations and survey. Boxes and whiskers indicate interquartile ranges for the data for each survey period, boxes are coloured according to month of year. Points show outliers. Daltons acoustic monitoring commenced in June 21 and was missing in January, February, most of March and November 2022. These months were excluded from analysis.

### Relationship of Pilbara Leaf-nosed Bat activity to environmental variables

#### 1. Chateau Cave in

Several GAMs were evaluated to assess the influence of environmental and temporal predictors on average nightly activity inside the main roosting chamber at Chateau Cave. Each model included an AR1 correlation structure, with estimated rho values between 0.56 and 0.62, indicating moderate temporal autocorrelation in nightly activity. This suggests that activity on a given night was related to activity on preceding nights. Including the AR1 structure allowed environmental predictors, including moon illumination and cumulative rainfall, to be assessed while accounting for short-term temporal dependence.

Model comparisons were based on ΔAICc values (Table 3). The base model, containing only DOY and year, explained 39.8% of deviance. Models that included environmental predictors improved model fit. The model with the lowest AICc and highest deviance explained (51.4%) included an interaction between moon illumination and cumulative rainfall, and artificial light at night (Table 3, Supplementary Table 1, Figure 3 and 4). Activity inside Chateau Cave declined when the light tower was operating, under higher moon illumination and with higher cumulative rainfall. The moon illumination × cumulative rainfall interaction predicted highest activity under low moon illumination and low cumulative rainfall, and lowest activity under high cumulative rainfall regardless of moon illumination. DOY and year were also significant, with activity increasing mid-year in June and July, remaining similar between 2021 and 2022, and declining markedly in 2023. Cave entry was associated with increased activity in one candidate model, but this effect had limited explanatory power and was not retained once stronger environmental predictors were included.

**Table 3.** Overview of models with explained deviance > 50% to assess the association between Chateau Cave in activity and environmental variables. Models are sorted according to the AIC value. All models including the base model included an AR1 model error structure and a cyclic cubic spline smooth for DOY and categorical variable year to assess temporal trends not explained by environmental variables. Model predictors in addition to the base variables are listed in the second column. The deviance explained indicates the proportion of the model’s total deviance accounted for by the model. The last column lists all smooth and categorical variables with P<0.050. Please refer to Table 1 for a detailed description of the predictors.

| Model name | Model Predictors in addition to s(DOY) and year (factor) | $\Delta AIC_c$ | Deviance explained | Predictors with significant effects ($P < 0.050$) |
| --- | --- | --- | --- | --- |
| **GAM.CCin.1** | moon x rain, artificial light | 0.0 | 51.4% | Interaction between cumulative rain and moon, artificial light, DOY, year |
| **GAM.CCin.2** | moon x rain | 45.8 | 50.0% | Interaction between cumulative rain and moon, DOY, year |
| **GAM.CCin.3** | rain, mean barometric pressure, artificial light | 53.4 | 49.9% | Cumulative rain, mean barometric pressure, artificial light DOY, year |
| **GAM.CCin.base** |  | 101.8 | 39.8% | DOY and year |

**Figure 3.**
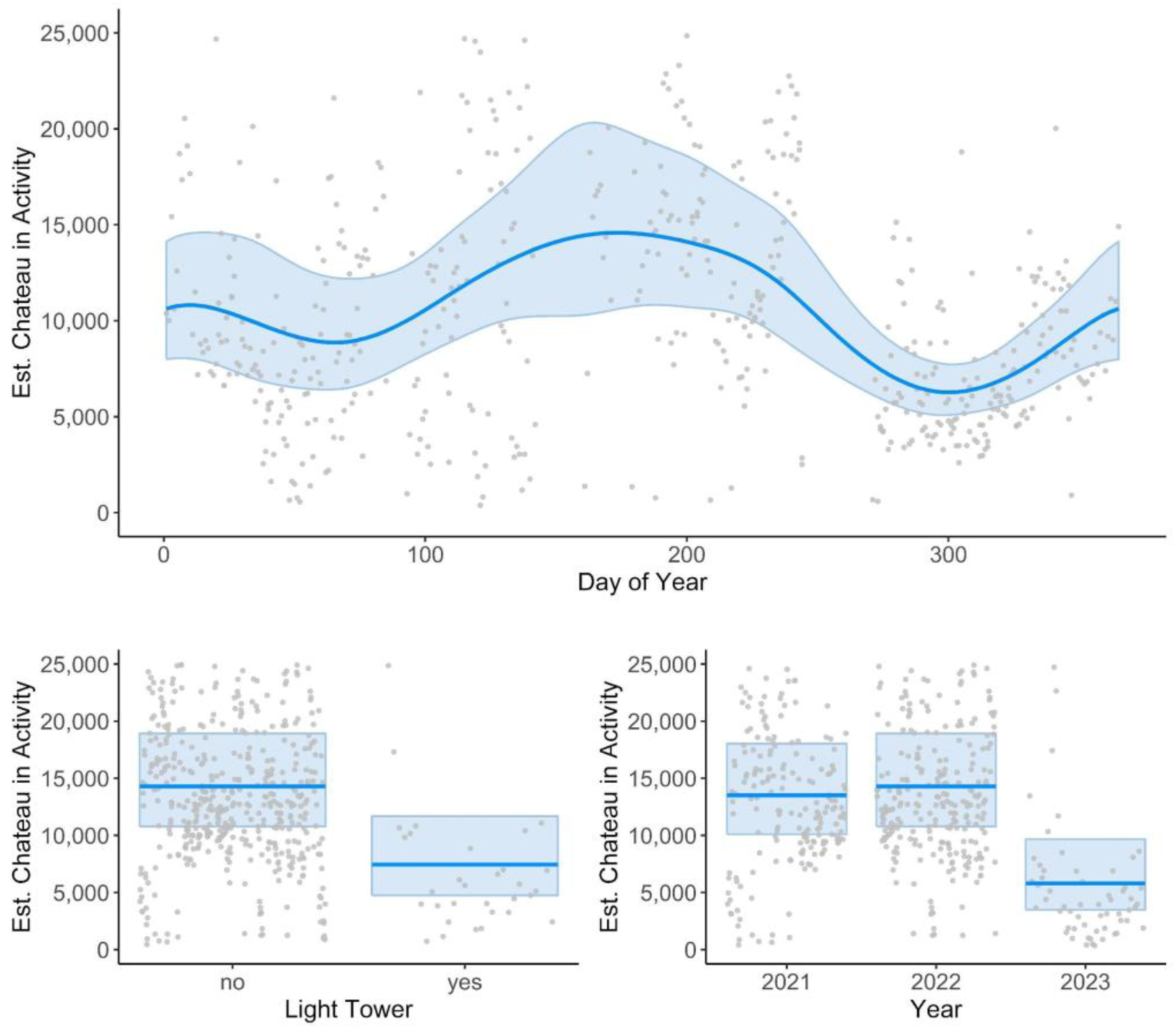
Estimated Chateau Cave in activity based on the best fit GAM m1 with A) predictors DOY, year and light tower operation (with other predictors held at their mean or reference level) and

**Figure 4.**
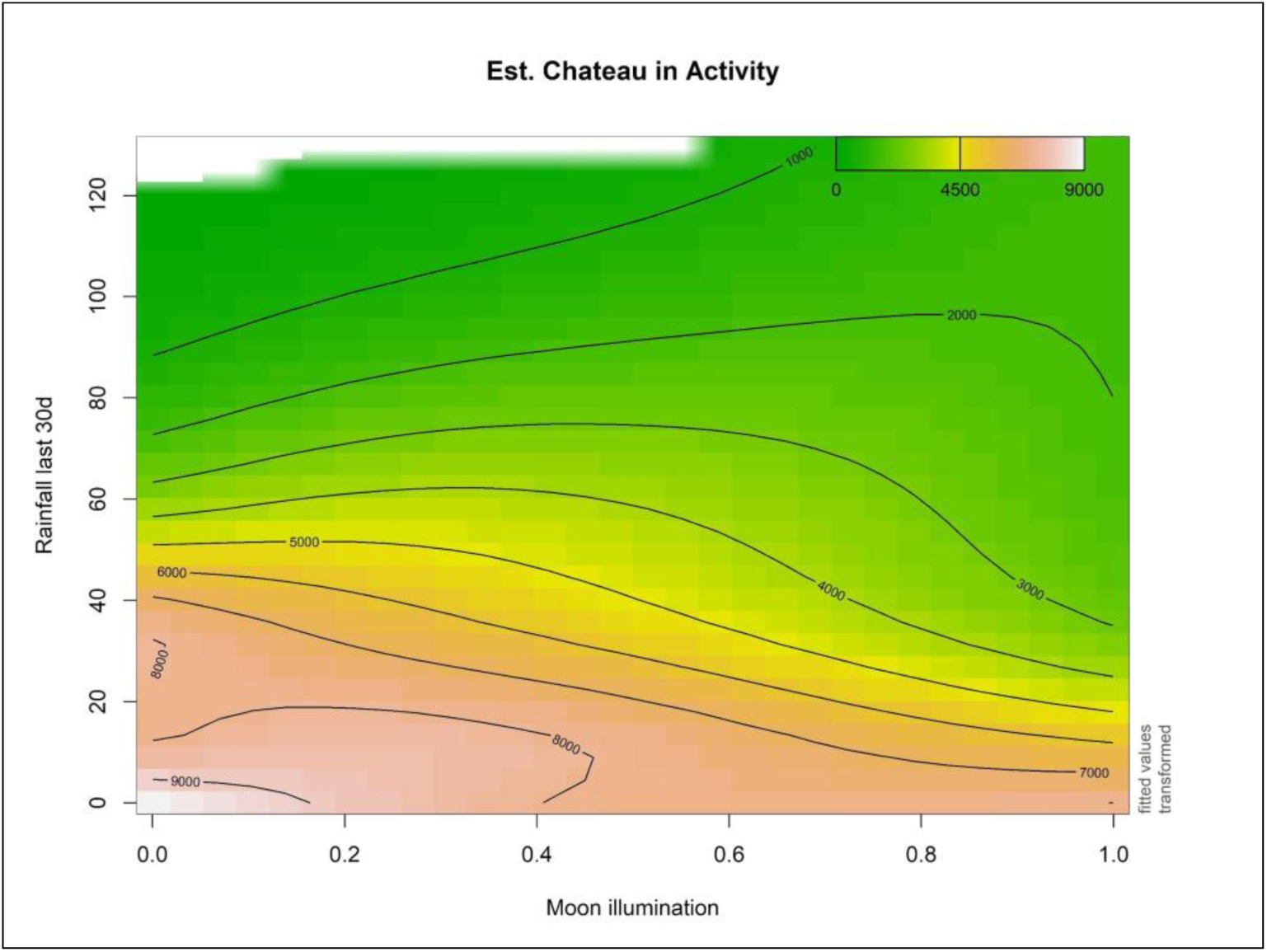
interaction of total rainfall of previous 30 days and moon illumination (with light tower “off”, year 2022 and DOY 1). Orange-red areas mark high Pilbara Leaf-nosed Bat activity and green low. The white area at the top left indicates a lack of available data for this combination of predictor values.

#### 2. Chateau Cave out

As with the internal detector at Chateau Cave, several GAMs were evaluated to assess environmental and disturbance-related predictors of nightly activity outside the cave entrance. AR1 rho estimates ranged from 0.48 to 0.59, indicating moderate temporal autocorrelation, although temporal dependence was slightly weaker than for the internal detector.

The best-supported model explained 61.9% of deviance and included a cumulative rainfall × moon illumination interaction, mean humidity and mean temperature (Table 4, Figure 5 and 6, Supplementary Table 2). Activity outside Chateau Cave was highest under low moon illumination and low cumulative rainfall. Mean temperature and mean humidity were also significant predictors. Compared with the internal detector, human disturbance variables were not strongly associated with activity outside the cave entrance. Light tower operation showed a negative trend, but this was not statistically significant. Year was significant, with activity lower in 2022 than 2021 and declining further in 2023.

**Table 4.**
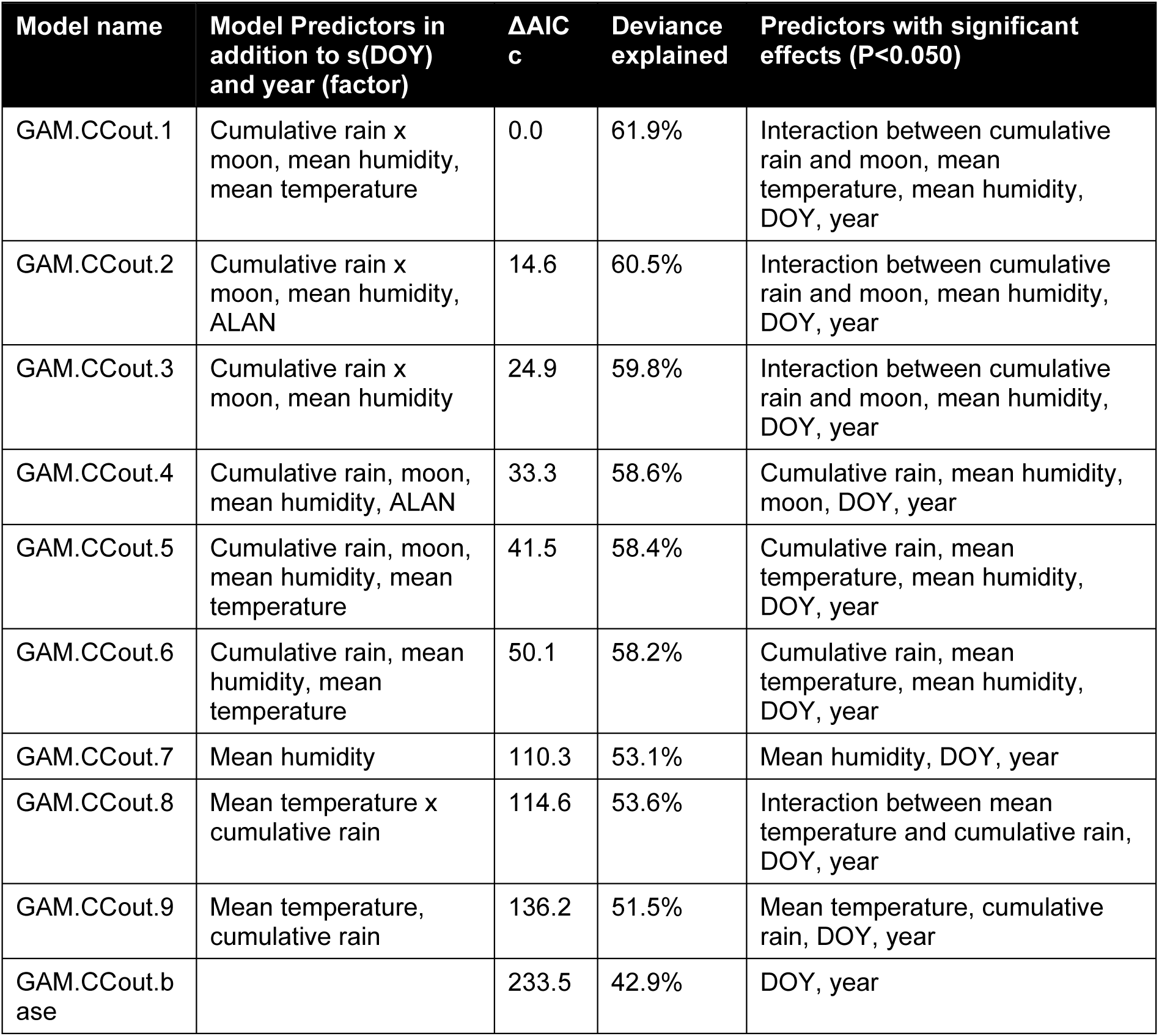
Overview of models with explained deviance > 50% to assess the association between Chateau Cave out activity and environmental variables. Models are sorted according to the AIC value. All models including the base model included an AR1 model error structure and a cyclic cubic spline smooth for DOY and categorical variable year to assess temporal trends not explained by environmental variables. Model predictors in addition to the base variables are listed in the second column. The last column lists all smooth and categorical variables with P<0.050. Please refer to Table 1 for a detailed description of the predictors.

**Figure 5.**
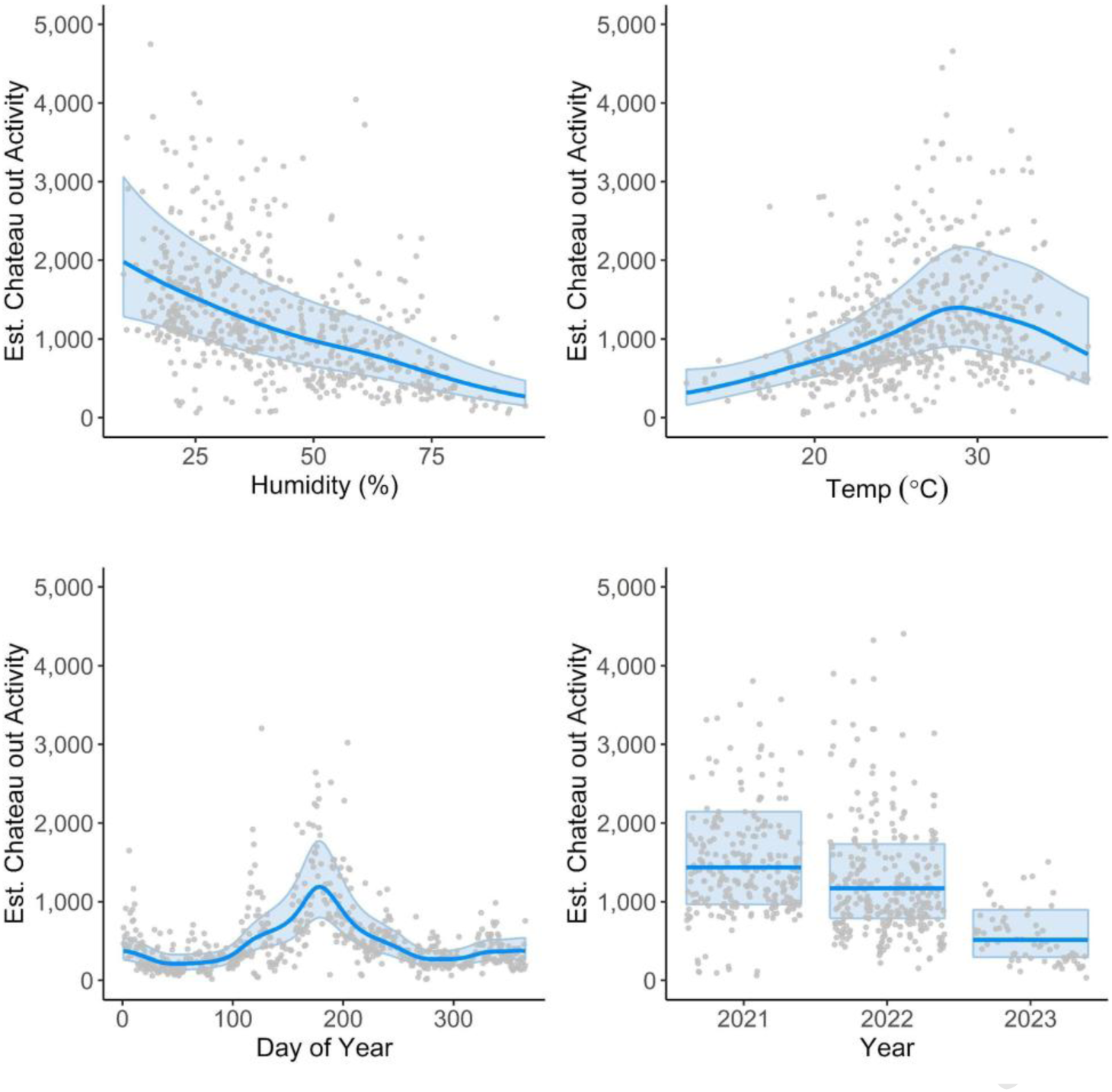
Estimated Chateau Cave out activity based on the best fit GAM.CCout.1 with predictors mean humidity, temperature, DOY and year (with other predictors held at their mean or reference level)

**Figure 6.**
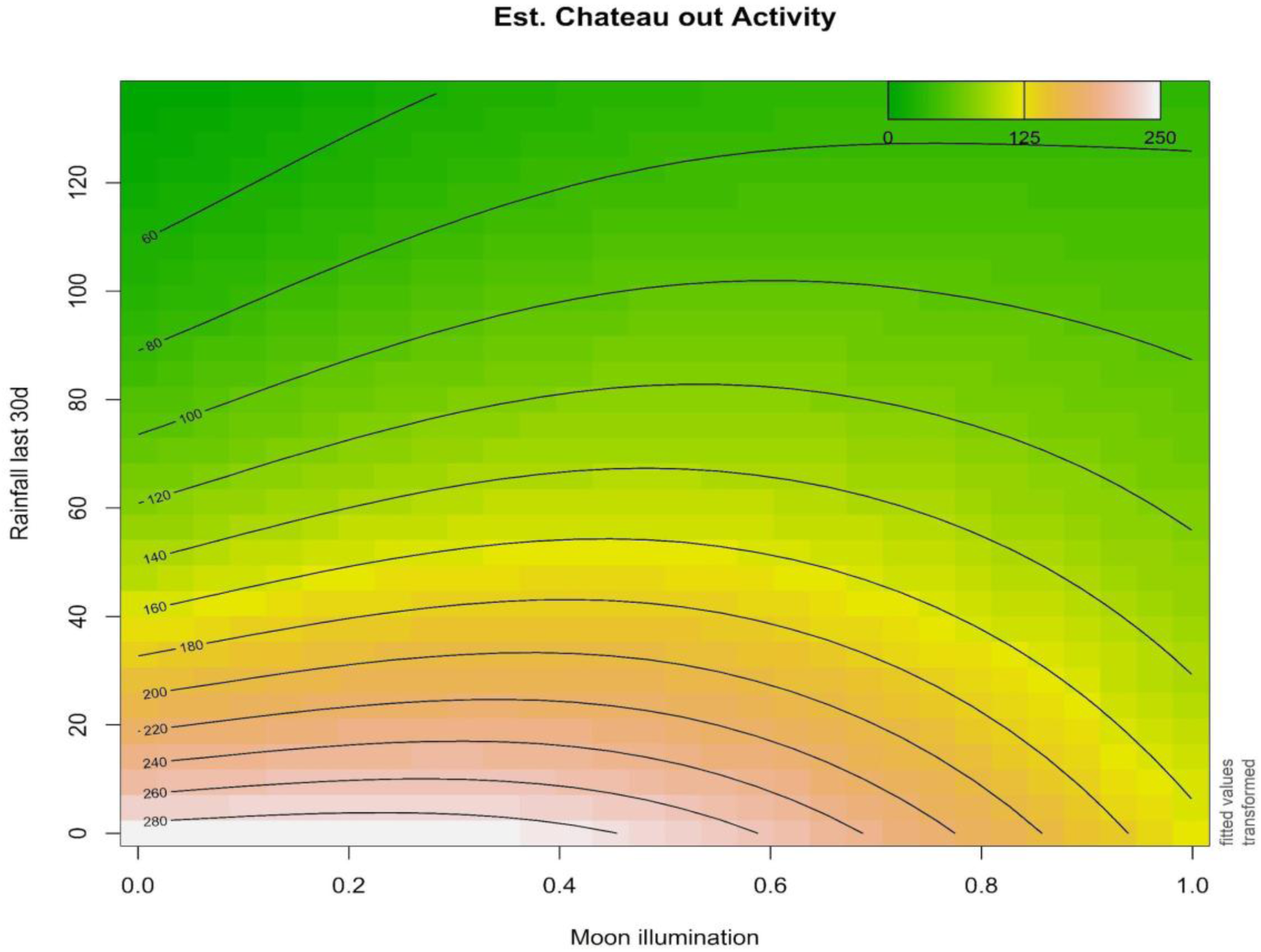
an interaction of total rainfall of previous 30 days and moon illumination (with year held at 2022, DOY 1, temperature 30°C and humidity 60%). Daltons

#### 3. Daltons

The best-supported Daltons model explained 47.3% of deviance and included moon illumination, cumulative rainfall and mean temperature (Table 5, Figure 7 and 8, Supplementary Table 3). The interaction between cumulative rainfall and moon illumination was significant. As at Chateau Cave, activity declined with higher moon illumination. In contrast to Chateau Cave, activity at Daltons was highest when cumulative rainfall was high and moon illumination was low. Mean temperature was also significant, with activity increasing with temperature to approximately 28 °C before declining at higher temperatures. Fire improved model fit in some candidate models but was not retained in the best-supported model. Year was significant, with activity lower in 2022 than 2021, but not significantly different between 2023 and 2021. GAMs were evaluated using the same general approach as for Chateau Cave because environmental variability, fire and potential disturbance at distant roosts could influence activity. AR1 rho estimates ranged from 0.55 to 0.72, indicating stronger temporal autocorrelation than observed at Chateau Cave.

**Table 5.** Overview of models with explained deviance > 40% to assess the association between Daltons Cave out activity and environmental variables. Models are sorted according to the AIC value. All models including the base model included an AR1 model error structure and a cyclic cubic spline smooth for DOY and categorical variable year to assess temporal trends not explained by environmental variables. Model predictors in addition to the base variables are listed in the second column. The last column lists all smooth and categorical variables with P<0.050. Please refer to Table 1 for a detailed description of the predictors.

| Model name | Model Predictors in addition to s(DOY) and year (factor) | $\Delta AIC_c$ | Deviance explained | Predictors with significant effects ($P < 0.050$) |
| --- | --- | --- | --- | --- |
| GAM.Dal.1 | Moon, cumulative rain, mean temperature | 0.0 | 47.3 | Interaction between cumulative rain and moon, mean temperature, DOY, year |
| GAM.Dal.2 | Mean temperature, moon | 7.5 | 42.1 | Mean temperature, moon, DOY, year |
| GAM.Dal.3 | Moon, mean humidity, mean temperature | 8.8 | 43.0 | Moon, mean temperature, DOY, year |
| GAM.Dal.4 | Moon, cumulative rain | 15.5 | 42.2 | Moon, DOY, year |
| GAM.Dal.5 | Fire, moon, cumulative rain | 16.4 | 42.9 | Moon |
| GAM.Dal.5 | Moon, cumulative rain | 18.1 | 40.3 | Interaction between cumulative rain and moon, DOY, year |
| GAM.Dal.6 | Mining, Fire, moon, cumulative rain | 18.3 | 40.7 | Moon |
| GAM.Dal.base |  | 56.0 | 15.3 | Year |

**Figure 7.**
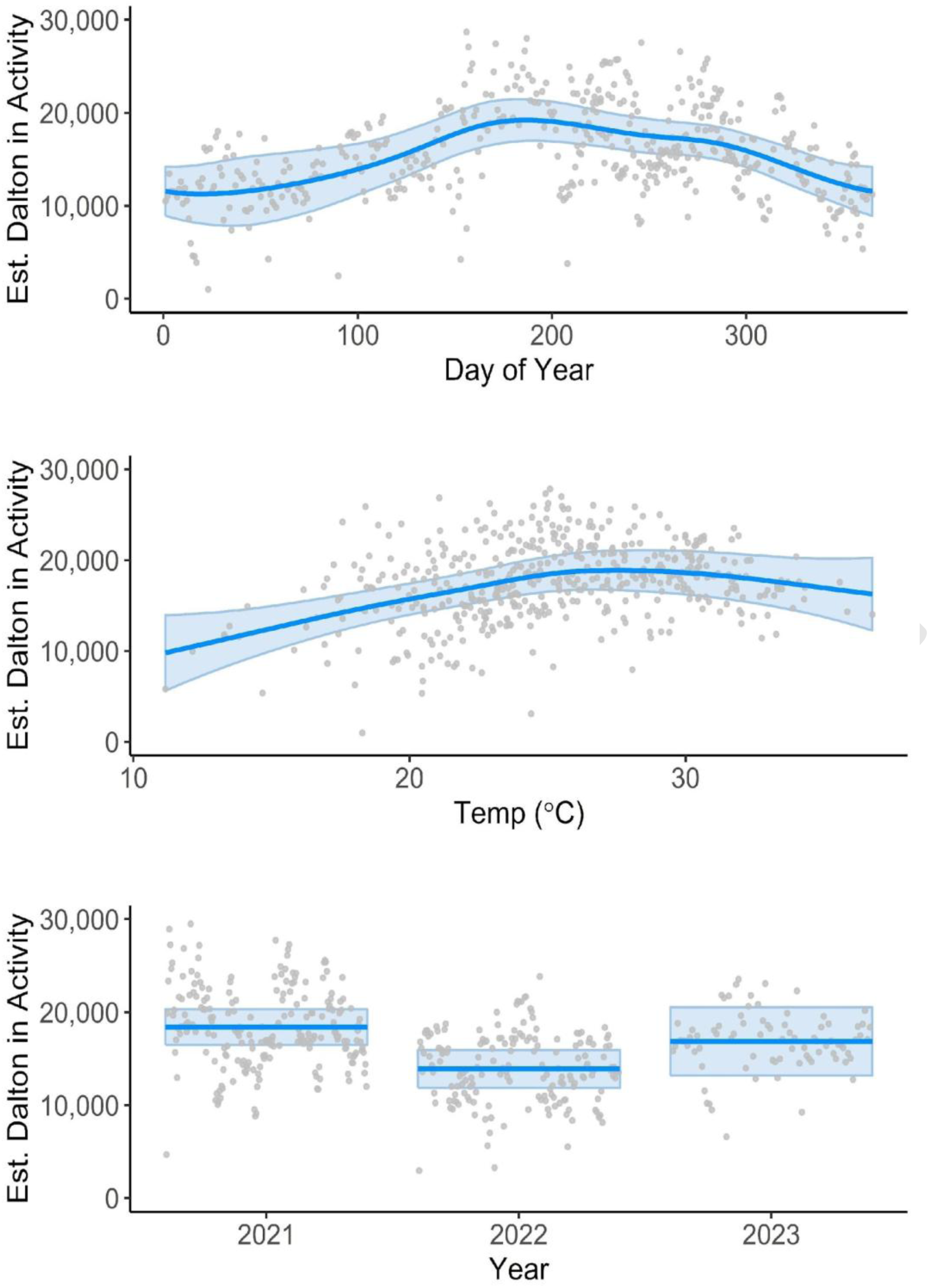
Estimated Daltons activity based on the best fit GAM.CCout.1 with predictors mean temperature, DOY and year with other predictors held at their mean or reference level) The blue band marks the 95% confidence interval and the dots are the partial residuals.

**Figure 8.**
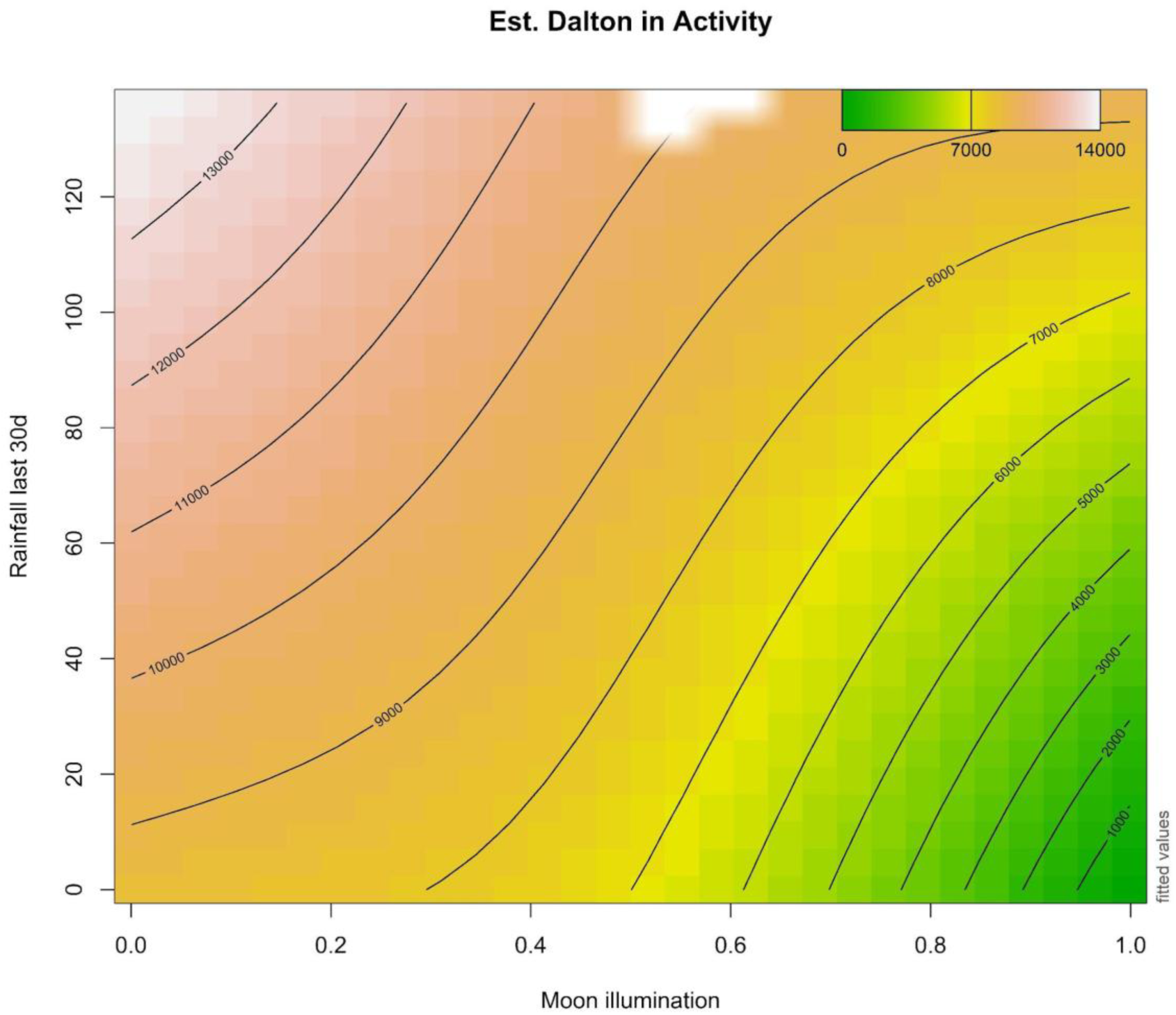
Interaction of total rainfall of previous 30 days and moon illumination (with year held at 2022, DOY 1, temperature 30°C and humidity 60%).

## Discussion

This study demonstrates substantial temporal variation in the activity of a small-bodied, long-lived and highly mobile mammal occupying an unpredictable arid environment. Unlike the rapid demographic responses to resource pulses characteristic of many small terrestrial arid-zone mammals (Ostfeld and Keesing 2000), bats are generally limited to producing one or two young per year (Racey and Entwistle 2000). Short-term variation in Pilbara Leaf-nosed Bat activity is therefore more likely to reflect behavioural responses than rapid demographic change. Activity at diurnal roosts was most strongly associated with the interaction between moon illumination and cumulative rainfall, although the direction and strength of these relationships differed among detector locations. Activity also differed markedly between detector positions at Chateau Cave. Together, these results demonstrate that acoustic activity at permanent diurnal roosts varies substantially with environmental conditions, time and detector location, with important implications for the interpretation of short-term acoustic surveys.

### Arid-zone variability and interpretation of acoustic activity

The contrasting relationship between cumulative rainfall and activity at Chateau Cave and Daltons indicates that rainfall does not operate as a simple regional driver of Pilbara Leaf-nosed Bat activity. At Chateau Cave, activity was highest under low moon illumination and low cumulative rainfall, whereas at Daltons activity was highest under low moon illumination and higher cumulative rainfall. These contrasting responses are consistent with the spatially heterogeneous nature of rainfall and subsequent productivity in arid landscapes, where similar climatic conditions do not necessarily produce equivalent ecological conditions among locations.

The contrasting rainfall responses may be mediated by local roost structure, surrounding foraging habitat, prey availability, predation risk and movement among roosts. Chateau Cave is a relatively deep, dark cave in banded ironstone and is known to maintain consistently high temperature and humidity (North et al. in prep). It may therefore function as a refuge during drier conditions. By contrast, Daltons is a narrow fissure within conglomerate rock. The internal roosting area is inaccessible, but the physical structure of the cave suggests that its microclimate may be more variable and more strongly influenced by external conditions. Rainfall may also influence invertebrate availability, although the relationship is likely to be taxon-specific and spatially variable. Invertebrate activity is often highest during pulses of productivity following rainfall (Westerhuis et al. 2020; Palmer 2010), although responses differ among taxa and landscapes (Kwok et al. 2016). We did not measure food availability, and the diet of Pilbara Leaf-nosed Bat in the Pilbara is not well documented, but the northern population of R. aurantia selectively eats moths and beetles (Churchill 1994).

Spatial responses of insectivorous bats to arid-zone conditions have been demonstrated elsewhere in Australia. In central Australia, differences in bat activity between riparian woodland and surrounding vegetation became more pronounced during hot, dry conditions, while activity also varied substantially among river systems (Westerhuis et al. 2021). This suggests that environmental conditions can alter the relative use of habitats across arid landscapes rather than producing uniform changes in activity across locations.

### Evidence for lunar avoidance in Pilbara Leaf-nosed Bat

While a relationship between rainfall and Pilbara Leaf-nosed Bat activity has previously been hypothesised (Bat Call WA 2021), this study provides evidence of a negative association between moon illumination and recorded Pilbara Leaf-nosed Bat activity. Activity was generally lower under higher moon illumination, consistent with lunar phobia or lunar avoidance reported in other bat species (Börk 2006; Zeppelini et al. 2019; Vasquez et al. 2020). In contrast to cohabiting cave bats such as the common sheath-tailed bat (*Taphozous georgianus*) and Finlayson’s cave bat (*Vespadelus finlaysoni*), Pilbara Leaf-nosed Bat emerge from cave roosts after civil twilight in full darkness. Avoidance of high-light conditions may reduce predation risk in open environments (Rydell and Speakman 1995), which is particularly relevant in the Pilbara where vegetation cover is naturally sparse.

Responses to moon illumination vary among bat species and environments. Species in tropical systems have been found to be more sensitive to moon illumination than temperate species (Saldaña-Vázquez and Munguía-Rosas 2013; Apoznański et al. 2024), and given the tropical origins of Pilbara Leaf-nosed Bat (Armstrong 2001), the observed negative relationship with moon illumination is plausible. The persistence of a moon illumination effect at Chateau Cave, where the entrance was also subject to artificial light at night, suggests that activity may respond to broader lunar-cycle cues or landscape-scale light conditions rather than only immediate illumination at the cave entrance.

### Artificial light and anthropogenic disturbance

Artificial light at night was associated with reduced activity inside Chateau Cave, but not outside the cave entrance. This result is consistent with the possibility that Pilbara Leaf-nosed Bat are sensitive to light spill near roosts. However, this inference should be treated cautiously because the light tower operated intermittently in late 2022 and continuously from early 2023, meaning that artificial light and year were not fully independent in the model. The marked decline in activity at Chateau Cave in 2023 may be related to continuous artificial light operation, but other unmeasured year-specific factors could also have contributed.

Artificial light at night has been shown to negatively affect some bat species, with responses often related to foraging behaviour and flight ecology. Fast-flying species that forage above canopy height may be less sensitive than species that forage within or near vegetation (Li et al. 2024; Frank et al. 2018). In contrast, *Rhinolophidae* species, a sister family to *Rhinonicteridae* within the *Rhinolophoidea*, are consistently light-avoidant (Luo et al. 2021; Straka et al. 2019).

Other anthropogenic disturbance variables had weaker support. Cave entry was associated with increased activity in one candidate model for Chateau Cave in, but this relationship was not retained once stronger environmental predictors were included. The study therefore provides no strong evidence that the cave-entry regime used here produced a measurable negative effect on nightly acoustic activity. Blasting and mining activity were also not retained as strong predictors in the best-supported models. These findings should not be interpreted as evidence that such activities are harmless, but rather that their effects were not clearly separable from environmental and temporal variation in the available dataset.

### Temperature, humidity and roost-specific responses

Aside from the interaction between cumulative rainfall and moon illumination, other environmental predictors had weaker and more detector location-specific relationships with Pilbara Leaf-nosed Bat activity than expected. Humidity was a significant predictor of reduced activity outside Chateau Cave, with very low activity recorded when humidity exceeded 75%, but this relationship was not observed at Daltons. High humidity can increase ultrasonic attenuation and reduce detection distance (Armstrong and Kerry 2011), potentially contributing to the lower activity recorded outside Chateau Cave at humidity above 75%. However, the absence of a comparable relationship at Daltons suggests that detectability alone may not explain this pattern. The observed relationship could reflect insect availability, flight conditions or physiological constraints, but further investigation is required.

Mean temperature was positively associated with activity at Daltons but not Chateau Cave. At Daltons, activity increased with mean daily temperature to approximately 28 °C and then declined at higher temperatures. This is consistent with the reported sensitivity of R. aurantia to temperatures outside the thermoneutral zone of approximately 30 ± 2 °C (Baudinette et al. 2000). During this study, mean daily temperatures ranged from 11 to 36 °C. Temperature is a common predictor of bat activity across a range of ecosystems, but there remains limited research on arid Australian bats. Given projected climate change in the Pilbara, further investigation of temperature effects on Pilbara Leaf-nosed Bat roost use and foraging activity is warranted.

### Fire, predation and disturbance interactions

Although fire was not significant in the best-supported Daltons model, some candidate models suggested reduced activity after fire. Increased bat activity after fire has been reported in some systems because of enhanced foraging conditions (Doty et al. 2016; Law et al. 2018), but little is known about how bats respond to fire in the Pilbara or in arid Australian landscapes more broadly. At cave roosts, the effects of fire may differ from effects in foraging habitat. Fire may alter vegetation structure around roost entrances, change prey availability or increase vulnerability to predators. In the months following the fire at Daltons, at least 76 Pilbara Leaf-nosed Bat were killed by a feral cat (Moyses et al. 2024). Feral cats are known to be less successful hunting in areas with mature spinifex (Triodia species) (Moore et al. 2024; Ngururrpa Rangers et al. 2024), which is a dominant vegetation type in the Pilbara. These observations suggest that interactions among fire, vegetation structure and predation risk should be a priority for future Pilbara Leaf-nosed Bat research.

### Roost networks and unexplained temporal variation

Despite significant relationships with moon illumination, cumulative rainfall, humidity, temperature and artificial light, a substantial proportion of temporal variation in Pilbara Leaf-nosed Bat activity remained unexplained. Spatial and temporal variation in activity has also been reported for other insectivorous bats in arid Australia. In central Australia, bat activity differed among habitats and river systems, and habitat associations changed through time, with activity in riparian woodland becoming more pronounced during hot and dry conditions (Westerhuis et al. 2021). Similarly, Aylen et al. (2025) found substantial variation in activity among detector locations and sampling nights, with spatial patterns for some species also changing through the night. Together with the temporally variable habitat associations reported by Westerhuis et al. (2021), these findings indicate that the distribution of insectivorous bat activity across arid landscapes is dynamic rather than spatially fixed.

For Pilbara Leaf-nosed Bat, extensive movement among permanent diurnal roosts provides a mechanism by which such spatial redistribution could occur over large areas. Individuals have been recorded moving up to 170 km between permanent diurnal roosts (Bullen and Reiffer 2019), undertaking nightly foraging movements of up to 40 km (Knuckey et al. 2024), and moving between roosts up to 60 km apart within a few hours (O’Brien and Westerhuis 2025). Frequent roost switching may simultaneously produce fission–fusion social dynamics, with individuals separating and reassociating among roosts through time (Kerth and König 1999). Whereas fission–fusion describes the resulting social organisation, facultative nomadism could provide an ecological explanation for broader shifts in spatial distribution.

Movement is an important behavioural response to resource limitation in arid-zone bats, including changes in foraging range, seasonal movements and shifts among alternative roosts (Conenna et al. 2025). Nomadism may represent an extension of this spatial flexibility where resources are particularly unpredictable in space and time. Nomadic movement is a widespread strategy among animals inhabiting unpredictable environments (Teitelbaum and Mueller 2019), allowing individuals to track changing resource distributions rather than remaining associated with fixed areas. We suggest that facultative nomadism could contribute to temporal variation in the use of permanent diurnal roosts by Pilbara Leaf-nosed Bat, particularly if individuals redistribute among roosts as environmental conditions and resource availability change across the landscape.

### Implications for acoustic surveys and impact assessment

The results presented here show that timing, duration and detector placement can influence conclusions drawn from ultrasonic surveys for Pilbara Leaf-nosed Bat. Although methods such as barrier sheeting, thermal counts of emerging bats, trapping, PIT tagging and radio tracking can provide complementary information on roost occupancy, colony size and individual movements, ultrasonic recording remains a widely used method for surveying and monitoring Pilbara Leaf-nosed Bat, particularly within environmental assessment and compliance programs (Bradley et al. 2024).

However, acoustic activity does not provide a direct measure of the number of individuals using a roost, and previous reviews have highlighted the need to validate acoustic-based estimates against video or other independent approaches (Bradley et al. 2024). For initial assessment of potential diurnal roosts, surveys should therefore incorporate temporal replication and, where practicable, sample across contrasting environmental conditions rather than relying on a single short deployment. At minimum, ultrasonic monitoring should span two separate survey periods selected to capture contrasting moon illumination and rainfall conditions. Where results will inform impact assessment, roost classification or decisions about habitat loss, longer deployments or complementary methods should be considered. Low activity during a short survey may reflect unfavourable lunar, rainfall or temperature conditions rather than low roost value.

These findings have implications beyond Pilbara Leaf-nosed Bat. Threatened bats in arid and semi-arid landscapes are often assessed under short survey windows driven by project timelines, access constraints and seasonal logistics. Our results show that such surveys should be interpreted cautiously unless they capture a meaningful range of environmental conditions. Where roost classification or impact assessment depends on acoustic activity, surveys should be temporally replicated, avoid reliance on single lunar or rainfall conditions, and incorporate relevant environmental covariates into interpretation. This is particularly important where low activity could be used to infer low roost significance.

More broadly, this study highlights a fundamental challenge in monitoring highly mobile species in climatically variable environments. Acoustic detectors sample activity at fixed locations, whereas the animals being monitored can respond to environmental variability through behavioural change and movement across much larger spatial scales. Long-term monitoring that incorporates environmental covariates and, where possible, simultaneous sampling across multiple sites will therefore provide a more robust basis for distinguishing local responses from broader changes in landscape use. For threatened cave-roosting bats in arid environments, understanding this spatial and temporal context is essential if acoustic activity is to be interpreted meaningfully in conservation and impact assessment.

## Acknowledgements

We gratefully acknowledge the Traditional Nymal Owners of the lands on which this study was undertaken and pay our respects to their Elders past, present, and emerging. We also recognise their deep connection to Country and their continuing role in caring for these lands. We gratefully acknowledge GHD Ecologists Emma DeMamiel, Lynette Grear, Robert Cooper Brown and Dylan Goldspink for assistance in the field. We also thank the Fortescue Metals Group Iron Bridge staff at North Star Mine for their invaluable logistical support during field work, particularly Sally Shepheard and Ziggy Nielson. We also extend our gratitude to Heliwest pilots for safely transporting personnel and our equipment to study detector locations. We thank Drew Farrer from GHD for his guidance and support in overseeing the project and for providing valuable input during its execution.

